# LncRNA-CIR6 Mediates Repair of Infarcted Hearts by Inducing MSCs Differentiation into Cardiogenic Cells through CDK1

**DOI:** 10.1101/2024.01.25.577303

**Authors:** Xiaotian Cui, Hui Dong, Shenghe Luo, Bingqi Zhuang, Yansheng Li, Chongning Zhong, Yuting Ma, Xianwu Cheng, Lan Hong

**Author notes:** Corresponding Author: Lan Hong, MD, PhD Associate Professor, Department of Physiology and Pathophysiology College of Medicine, Yanbian University, Yanji 133002, China.; Xianwu Cheng, MD, PhD Professor, Department of Cardiology and Hypertension Yanbian University Hospital, Yanji 133002, China. Xiaotian Cui and Hui Dong contributed equally to this work. The study is free from any potential conflicts of interest, both financial and non-financial.

## Abstract

**Purpose:** This study aims to investigate the induction effect of LncRNA-CIR6 on MSCs differentiation into Cardiogenic Cells *in vitro* and *in vivo*.

**Methods:** In addition to pretreatment with the Ro-3306 (CDK1 inhibitor), LncRNA-CIR6 was transfected into BMSCs and hUCMSCs using jetPRIME. LncRNA-CIR6 was transfected into C57BL/6 mice heart by 100 μL of AAV9-cTnT-LncRNA-CIR6-ZsGreen i.v. After 3 weeks of transfection followed by AMI surgery, hUCMSCs (5×10^5^/100 μL) were injected by i.v 1 week later. Cardiac function was evaluated using VEVO 2100 and electric mapping 9 days after cell injection. IF, Evans blue-TTC, Masson staining, FACS, and WB were used to determine relevant indicators.

**Results:** LncRNA-CIR6 induced a significant percentage of differentiation in BMSCs (83.00±0.58)% and hUCMSCs (95.43±2.13)% into cardiogenic cells, as determined by the expression of cTnT. Compared with MI group, cardiac contraction and conduction function in MI heart treated by LncRNA-CIR6 or combined with MSCs injection groups were significantly increased as well as the area of MI and fibrosis were significantly lower. The transcriptional expression region of LncRNA-CIR6 was in Chr17 from 80209290 to 80209536. The functional region of LncRNA-CIR6 was located at nucleotides 0-50/190-255 in the sequence. CDK1 is a protein found to be related to the proliferation and differentiation of cardiomyocytes is which located in the functional region of LncRNA-CIR6 secondary structure (from 0 to 17). Ro-3306 impeded the differentiation of MSCs into cardiogenic cells, while MSCs transfected with LncRNA-CIR6 showed high expression of CDK1. LncRNA-CIR6 mediates repair of infarcted hearts by inducing MSCs differentiation into cardiogenic cells through CDK1.

**Conclusions:** LncRNA-CIR6 mediates repair of infarcted hearts by inducing MSCs differentiation into cardiogenic cells through CDK1.

## 1. Introdction

Acute myocardial infarction (AMI) is a serious cardiovascular disease [1]. The proliferation of cardiomyocytes during normal aging cannot reverse the damage caused by acute myocardial infarction, and it is difficult for cardiomyocytes to regenerate, which will lead to abnormal recovery of cardiac function [2, 3]. However, cell therapy using stem cells can effectively compensate for the loss of cardiac myocytes, thus overcoming the difficulties of traditional medical methods [4].

Embryonic stem cells (ESCs) and induced pluripotent stem cells (iPSCs) have been used in clinical trials for cell therapies such as cristae injury, macular degeneration, and Type 1 diabetes [5, 6, 7]. However, even though the research on ESCs and iPSCs has made some progress, it still faces many technical challenges in practical application [8]. For example, how to ensure the stability and safety of cells, how to avoid immune rejection, and how to improve the efficiency of cell differentiation. Compared with iPSCs and ESCs, mesenchymal stem cells (MSCs), as the seed cells of regenerative medicine and tissue engineering, have many advantages, such as low immune rejection, easy preparation, large size of MSCs and no ethical influence [8, 9]. Studies have found that MSCs can promote myocardial regeneration after myocardial infarction [10]. Transplanting MSCs not only makes up for lost heart cells but also regulates immune factors [11]. Cardiac MSCs reduce inflammation and ameliorate myocardial infarction in rats with ischemia-reperfusion injury [12]. The MSCs and their secreted products decrease the gene expression of pro-inflammatory cytokines and interleukin in cardiomyocytes, while the expression of antioxidant gene superoxide dismutase is upregulated [13, 14]. Therefore, how to regulate the proliferation and differentiation of MSCs and how to use Mesenchymal stem cells to cure diseases are urgent needs of regenerative science. The study of long non-coding RNA (LncRNA) in stem cells is an active and fruitful field [15]. In recent years, more and more studies have shown that lncRNA plays an important role in the self-renewal, differentiation, reprogramming, and other processes of stem cells [15, 16, 17]. LncRNA, as an intracellular gene regulator, can mediate the development of mesenchymal stem cells [16, 17]. As LncRNA H19 can repair damaged myocardium by improving the viability of MSCs and the angiogenic capacity of endothelial cells [18]. LncRNA TCF7 may maintain the cellular viability and stem cell viability of MSCs by activating the Wnt pathway [19].

lncRNA-CIR6 from the human heart was first identified by the Center for Regenerative Medicine at Texas A & M University in Cormus, non-cardiomyocytes, such as mouse embryonic stem cells, mouse and human ips cells, and mouse embryonic fibroblasts, can be induced to differentiate into cardiomyocytes expressing cTnT *in vitro*. However, it is not clear whether lncRNA-CIR6 can also induce MSCs.

Therefore, we investigated the effects of lncRN-CIR6 on the induction of MSCs into cardiogenic cells *in vivo* and *in vitro*, and whether these effects can resist myocardial infarction damage *in vivo*.

## 2. Materials and Methods

### 2.1 Animals

Sprague Dawley(SD) rats (100 ± 5 g) and C57BL/6 mice (20 ± 1 g) were used, provided by the laboratory animal center of Yanbian University (License No. SYXK (Ji) 2020-0009). All male animals were kept in a clean, well-ventilated, and well-lighted environment with a temperature of 25-26℃ and a humidity of 70%. All animal experiments in this study were approved by the institutional animal protection and use committee of Yanbian University (approval No. YD20240124001), and followed the National Institutes of Health Guidelines for experimental animal protection.

### 2.2 Isolation and Culture of Rat Bone Marrow Mesenchymal Stem Cells (BMSCs)

SD rats were sacrificed by cervical dislocation and then immersed in 75% ethanol for 5 minutes. Rats were placed in a supine position on a super-clean bench, and bilateral femurs were removed under sterile conditions. The muscle tissue attached to the surface of the femur was removed cleanly (note to avoid damaging the epiphyseal), and the femur was placed in a sterile PBS-filled petri dish for later use. Subsequently, the ends of the epiphysis were cut to expose the medullary cavity. Use a syringe to aspirate the α-MEM complete medium (Shanghai XP Biomed Ltd., China) to flush the bone marrow cavity into a centrifuge tube, and swirl the cell suspension to disperse the cells.

Centrifuge at 1000 rpm for 5 minutes at room temperature. Discard the supernatant, resuspend in 10 ml complete medium to make a single-cell suspension, transfer to a 10cm culture dish, gently shake to mix, and place in a cell incubator at 37℃, 5% CO_2_, and saturated humidity. After 24 hours, replace the α-MEM complete medium, and then replace the liquid every 2 days. When the fusion rate reaches 80% to 90%, proceed with subculture.

### 2.3 Transfect LncRNA-CIR6 into MSCs *In Vitro*

Human umbilical cord mesenchymal stem cells (hUCMSCs) were purchased from Shanghai Kanglang Biotechnology Co., Ltd. BMSCs were isolated from the femur of SD rats. The plasmid containing the LncRNA-CIR6 sequence was constructed by Hanbio Biotechnology Co.Ltd, China. When the primary cells were cultured for 7 days and the cell confluence reached 70%-80%, JetPRIME DNA & siRNA (Ployplus, USA) transfection reagents were used to transfect the MSCs. After 9 days of transfection, immunofluorescence and flow cytometry were performed on the 10th day (Fig. 1).

**Figure 1.**
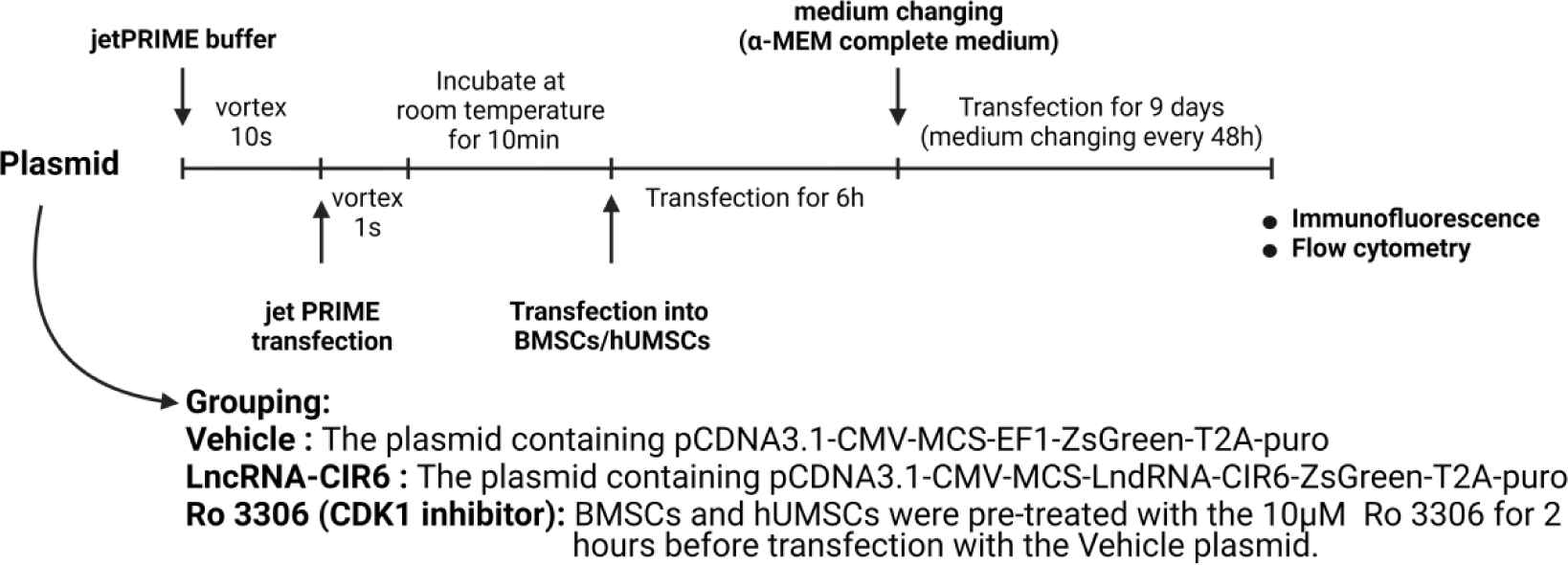
Experimental Protocols and Treatment Reagents *In Vitro*.

### 2.4 Experiment of Immunofluorescence

MSCs were divided into Vehicle, LncRNA-CIR6, and Ro 3306 groups according to the experimental protocol (Fig. 1). After discarding the old medium, transfected cells were fixed with 2 ml of 4% paraformaldehyde per well (10 min) and then removed. Subsequently, cells underwent three 5-minute washes with ice-based PBS (1 ml per well). Following permeabilization with 0.1% PBS-Tween, an overnight incubation with the primary antibody at 4℃, and the secondary antibody was conducted at room temperature for 1 hour away from light. After the washing steps, 10 μl of anti-fluorescence quenching sealing solution (Beyotime, China) containing DAPI was added for coverslipping. Finally, fluorescence was observed using a fluorescence inversion microscope (Thermo Fisher Scientific, USA). Primary antibody: Anti-Cardiac Troponin T antibody [EPR20266] at a dilution of 1:500 (Abcam, USA); Secondary antibody: Goat Anti-Rabbit IgG H&L(Alexa Fluor® 647)preadsorbed (ab150083) at a dilution of 1:2000 (Abcam, USA); Ro 3306: CDK1 inhibitor (APExBIO, USA)

### 2.5 Flow Cytometry

MSCs were divided into LncRNA-CIR6 and Vehicle groups for transfection (Fig. 1). On the 10th-day post-transfection, cell suspensions were collected (Density: 3 × 10^5^ cells/ml) and centrifuged at room temperature for 5 minutes in a 15ml centrifuge tube. The supernatant was discarded, and cells were fixed with 4% paraformaldehyde for 15 minutes to prevent cell cross-linking. After centrifugation, the supernatant was discarded, and cells were resuspended in pre-chilled 90% methanol for 15 minutes, facilitating further cell fixation and transparency treatment. Following two washes with PBS centrifugation, cells were resuspended in diluted primary antibody and incubated at 4℃ for 1 hour. Subsequently, cells were washed twice with PBS centrifugation and resuspended in diluted secondary antibody, followed by a 30-minute incubation at 4℃. After two additional washes, cells were resuspended in PBS and analyzed for all groups on a flow cytometer (Bdbiosciences, USA). Primary antibody: Anti-Cardiac Troponin T antibody [EPR20266] at a dilution of 1:500 (Abcam, USA); Secondary antibody: Goat Anti-Rabbit IgG H&L(Alexa Fluor® 647)preadsorbed (ab150083) at a dilution of 1:2000 (Abcam, USA).

### 2.6 Western Blot

MSCs were transfected according to the LncRNA-CIR6 group and Vehicle group, as illustrated in Figure 1. On the tenth day after transfection, the protein was extracted from each group of MSCs, and the protein concentration was determined using the BCA protein concentration assay kit (Solarbio, China). After quantification, the samples were mixed with an appropriate amount of protein tracer loading buffer (Proteintech, China) and denatured. The proteins were separated by SDS-PAGE gel electrophoresis and then transferred to the PVDF membrane (Merck, USA).

SDS-PAGE gel percentages were 12%, and blotting membranes were blocked with 5% skim milk for 1 hour. Subsequently, the membranes were incubated with the corresponding primary antibodies overnight at 4 °C. Afterward, the membranes were incubated with the corresponding secondary antibody for 1 hour at room temperature. Following exposure with the enhanced chemiluminescence kit (Merck, USA), the bands were processed using Image J software (National Institutes of Health, USA). Primary antibodies: Anti-CDK1 (phospho T161) + CDK2/CDK3 (phospho T160) antibody [EPR19546] (ab201008) at a dilution of 1:2000 (Abcam, USA) and β-Actin Antibody (4967S) at a dilution of 1:1000 (CST, USA); Secondary antibody: Anti-rabbit IgG, HRP-linked Antibody (7074S) at a dilution of 1:1000, (CST, USA).

### 2.7 Preparation of Adeno-associated Virus

The recombinant plasmid AAV9-cTnT-LncRNA-CIR6-ZsGreen (Hanbio Biotechnology Co.Ltd, China) was constructed by cloning the sequence of LncRNA-CIR6, green fluorescent protein ZsGreen, and cTnT promoter into the AAV9 vector. The AAV9-cTnT-LncRNA-CIR6-ZsGreen recombinant plasmid was transfected into HEK293T or other suitable cell lines, while the AAV9 helper plasmid and packaging plasmid were added to facilitate adeno-associated virus production and packaging. The AAV9-cTnT-LncRNA-CIR6-ZsGreen virus was purified and concentrated by centrifugation.

### 2.8 Experimental Protocols and Treatment Reagents *In Vivo*

The mice were randomly assigned to four groups: Sham group, MI group, LncRNA-CIR6 group, and LncRNA-CIR6 + huCMSCs group (Fig. 2). Sham group: Each mouse was injected with 100 μl of AAV9-cTnT-ZsGreen. After 3 weeks, a thoracotomy was performed without ligation, followed by the injection of 200 μl of normal saline one week later. MI group: Each mouse received an injection of 100 μl of AAV9-cTnT-ZsGreen. After a 3-week period, a thoracotomy was performed to induce MI, followed by the injection of 200 μl of normal saline one week later. LncRNA-CIR6 group: Each mouse was injected with 100 μl of AAV9-cTnT-LncRNA-CIR6-ZsGreen. After 3 weeks, a thoracotomy was performed to induce MI, followed by the injection of 200 μl of normal saline one week later. LncRNA-CIR6+hUCMSCs group: Each mouse was injected with 100 μl of AAV9-cTnT-LncRNA-CIR6-ZsGreen. After 3 weeks, a thoracotomy was performed to induce MI, and one week later, 200 μl of huCMSCs (injection volume: 5×10^5^ cells/100 μl) was administered.

**Figure 2.**
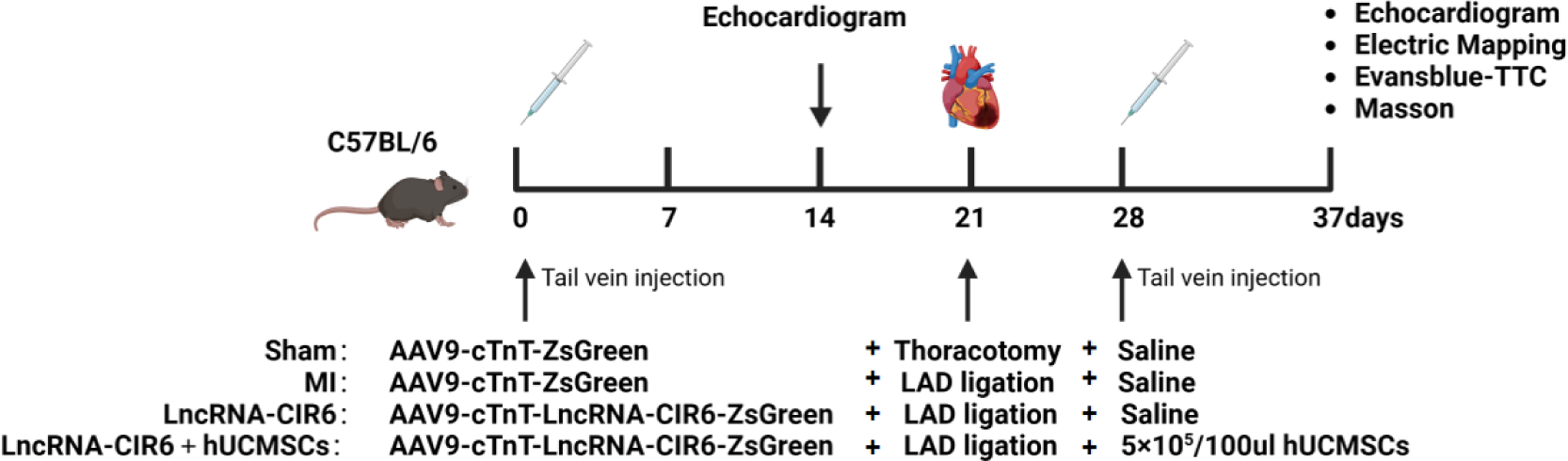
Experimental Protocols and Treatment Reagents *In Vivo*.

#### 2.8.1 Echocardiogram

Not only were echocardiograms performed on mice in each group one week before surgery, but also, after administering hUCMSCs or saline, observations were made for 9 days, followed by echocardiographic examinations of the mice’s hearts on the 10th day (Fig.2). The anesthetized mice were fixed on the operating platform in a supine position and maintained with isoflurane at a concentration of 2% and a flow rate of 0.6 L/min using an anesthetic mask. The mouse’s paw is attached to the electrode pad. Ensure that correct cardiac ultrasound is collected, mouse body temperature is maintained, and respiratory rates are monitored for physiological assessment and monitoring during imaging. Hair removal was performed on the abdomen and chest, and a thick layer of ultrasonic coupler was applied to the left side of the mouse’s chest. The anterior and posterior wall thickness at the maximum diameter of the long/short axis section, left ventricular internal dimension in systole, left ventricular internal dimension in diastole, left ventricular ejection fraction, and left ventricular fractional shortening were obtained by measurement. The instrument is the VEVO 2100 imaging system (American VisualSonics company, USA).

#### 2.8.2 Electric Mapping

Anesthetized mice in each group underwent thoracotomy, exposing the heart. A multi-channel electrode was attached to the free wall of the left ventricle. The multi-electrode array (MEA) consists of 32 electrodes with a diameter of 0.1 mm, in a 6×6 grid (size: 6 mm x 6 mm), The electrode spacing is 1.2 mm. 32 recording electrodes are connected to a 32-channel amplifier and data acquisition system (EMS64-USB-1002, MappingLab LTD., UK). The sampling frequency of each channel was set to 10kHz, and the field potential recording corresponded to that of the reference electrode placed on the cardiac perfusion metal cannula. MEA used EMapScope (MappingLab Limited, UK) software to provide 32 channels of non-invasive synchronous recording of cell field potential. The EMapScope software allows real-time monitoring of field potentials across all 32 channels, based on which heart rate can be calculated. The activation time of each channel is calculated based on the relative delay of the first detected waveform in the recording array. In order to generate an epicardial propagation map, the activation time of each channel is represented by a color code in a 6×6 grid that displays the original recording array to generate an epicardial propagation map for a clearer view of the conduction pattern and direction.

#### 2.8.3 Fluorescence and Masson Staining of Frozen Sections

After the completion of the final echocardiogram, the hearts of mice in each group were cut along the plane of ligation, with the ligation point serving as the horizontal reference. The cardiac tissues below the ligation point were preserved, frozen in OCT compound, and sectioned using a microtome to obtain 10 μm-thick slices. A portion of these slices was utilized for observing frozen cardiac tissues of mice in each group under a fluorescence microscope (Thermo Fisher Scientific, USA). Another set of slices was subjected to Masson staining. The frozen cardiac tissue slices from each group were stained following Masson’s Trichrome Staining Kit (Solarbio, China).

Subsequently, the degree of fibrosis was assessed using an inverted microscope (Thermo Fisher Scientific, USA).

#### 2.8.4 Evans Blue-TTC Staining

After administering hUCMSCs or saline to mice in each group, a 9-day observation period followed. On the 10th day, mice were anesthetized, and their chests were opened. The ascending aorta was carefully clamped with hemostatic forceps, and 0.5% Evans blue dye (Solarbio, China) was slowly injected from the proximal end of the clamp. Rapid bluing of the mouse heart was observed, with no significant changes noted below the ligature line. The heart was then extracted, swiftly cleaned in PBS, frozen at −80℃, cut into 0.5 mm slices, dyed in a 2% TTC solution (Solarbio, China) at 37℃ for 15 minutes, and photographed.

### 2.9 LncRNA-CIR6 Bioinformatics and Statistical Analysis

Analyze the possible related functions of LncRNA-CIR6 using Annolnc2 (http://annolnc.gao-lab.org/), such as its expression, subcellular localization, interacting miRNAs, or transcription factors that regulate it.

### 2.10 Statistical Analysis

Experimental data were statistically processed using GraphPad Prism 9.0 software. All data are presented as means ± S.E.M. Comparisons between two independent samples were performed using an unpaired t-test. Comparisons between multiple groups were conducted using one-way ANOVA and two-way ANOVA, with a significance level set at *P*<0.05.

## 3. Results

### 3.1 Effects of LncRNA-CIR6 on the Transformation of BMSCs and hUCMSCs into Cardiogenic Cells

To verify whether LncRNA-CIR6 can transform BMSCs and hUCMSCs into cardiogenic cells, a myocardial-specific marker, cTNT antibody, was selected for observation and analysis under fluorescence microscopy. Furthermore, the fluorescence intensity of APC was detected by flow cytometry to evaluate the efficiency of LncRNA-CIR6 in inducing the transformation of MSCs into cardiogenic cells. The results showed that on the 10th day after transfection with LncRNA-CIR6, green fluorescence was expressed in the cytoplasm of both MSCs, indicating successful transfection, and these cells also clearly expressed the red fluorescently labeled cTnT protein. In contrast, BMSCs and hUCMSCs transfected with empty plasmids only exhibited green fluorescence (Fig. 3A-B). This suggests that LncRNA-CIR6 can induce the differentiation of both MSCs into cardiogenic cells. The average conversion rate of LncRNA-CIR6 to cardiogenic cells in BMSCs was (83.00±0.5774)% (Fig. 3C-D), and in hUCMSCs, it was (95.43±2.1300)% (Fig.3E-F).

**Figure 3.**
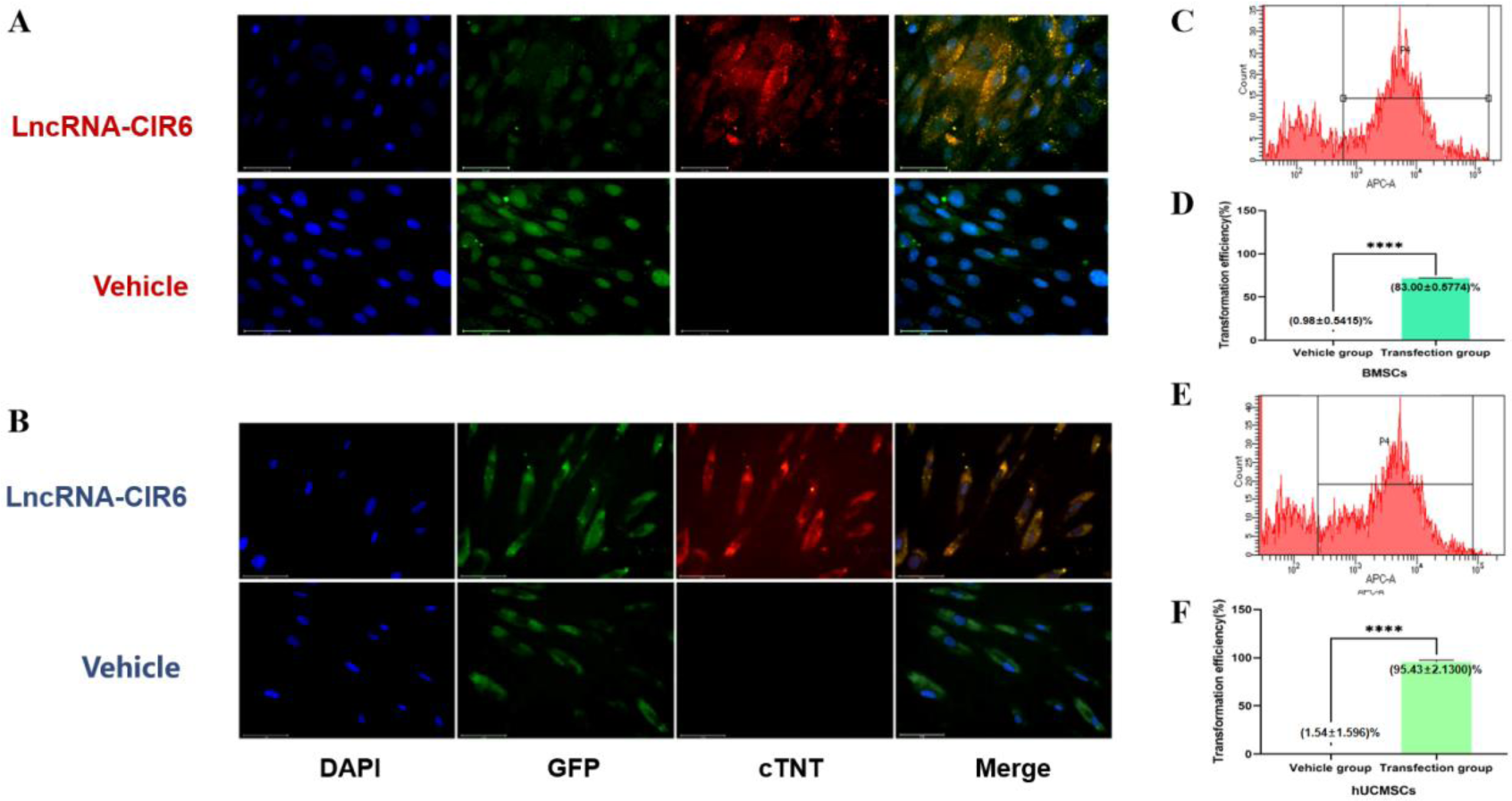
Effects of LncRNA-CIR6 on the Transformation of BMSCs (A, C-D) and hUCMSCs (B, E-F) cardiogenic cells. Scale: 20 μm (A-B); \*\*\*\**P*<0.0001 vs Vehicle group (C-F). (means ± SE, n = 3)

### 3.2 The Effects of LncRNA-CIR6 or Combined hUMSCs on Heart Function and Cardioprotection *In Vivo* in Mice with Myocardial Infarction

To assess contractile and conduction functions, we employed echocardiogram and electric mapping, respectively. The animal ultrasound system was used to compare cardiac function before and after MI, with the M-mode echocardiogram presented in Fig. 4A. Prior to the operation, the left ventricular contraction function was robust, and the left ventricular wall exhibited normal characteristics. After the operation, the anterior wall of the left ventricle displayed thinning, weakened or disappeared segmental motion, impaired contraction function, and tissue remodeling. Importantly, cardiac function in both the LncRNA-CIR6 group and the LncRNA-CIR6 + hUCMSCs group surpassed that of the MI group.

**Figure 4.**
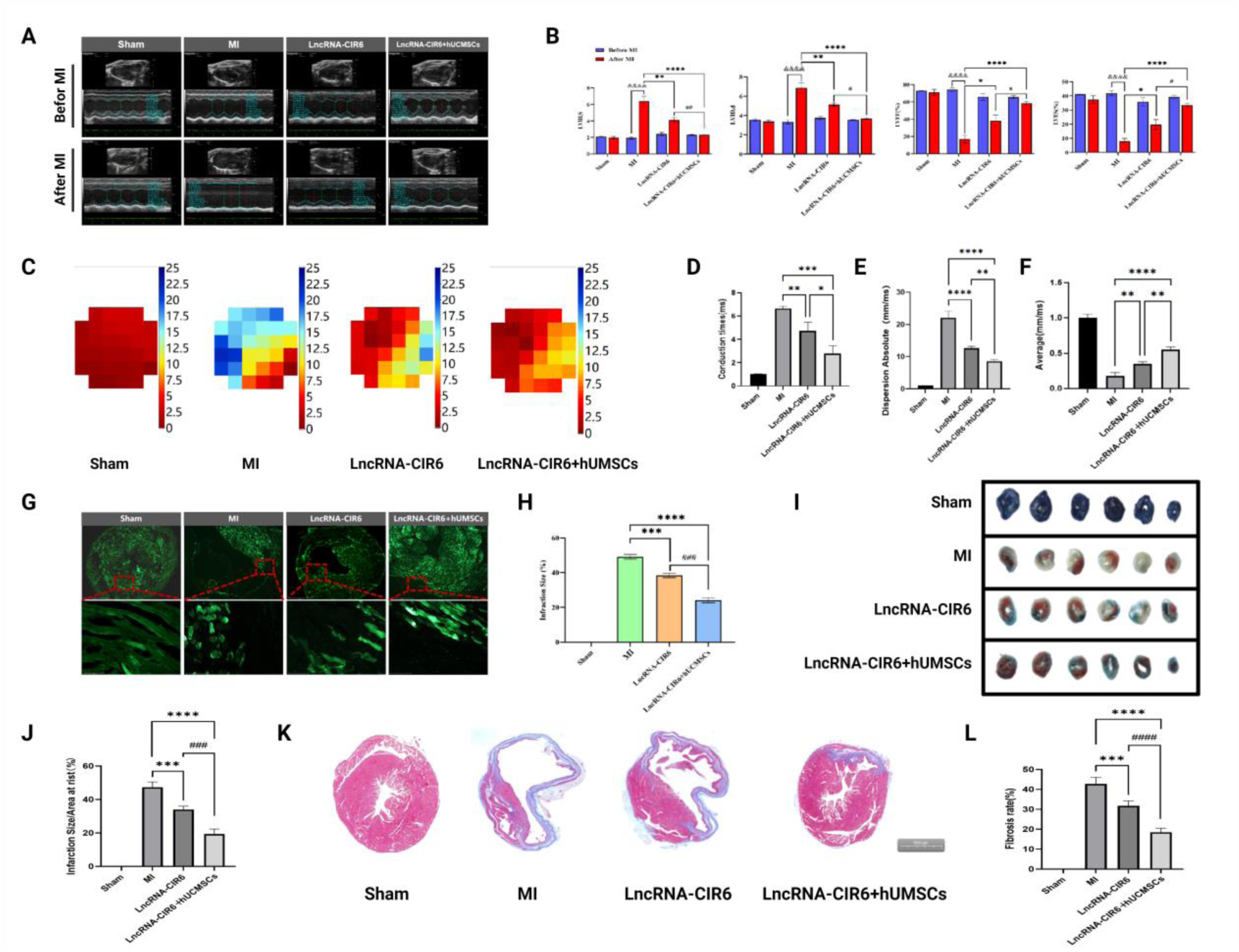
The Effects of LncRNA-CIR6 or Combined hUMSCs on Heart Function and Cardioprotection *In Vivo* in Mice with Myocardial Infarction. A: Echocardiogram. B: Cardiac Parameters Measured: LVIDs (Left ventricular internal dimension in systole); LVIDd (Left ventricular internal dimension in diastole); LVEF (Left ventricular ejection fraction); LVFS (Left ventricular fractional shortening). C: Ventricular Conduction Time Diagram: Visual representation of ventricular conduction time diagram (Fig. 4C). Illustrates conduction time from fast to slow, where the transition from red to blue signifies increasing conduction time. More red indicates faster conduction. D: Visual Statistical Representation of Fig. 4C. E: Dispersion Absolute of Ventricular Conduction: Represents the heterogeneity of ventricular conduction; a smaller dispersion absolute suggests improved heart function. F: Average Ventricular Conduction Velocity. G: Fluorescence Imaging of Frozen Mouse Heart Slices (Scale: 2 mm; 200 μm): Bright fluorescence indicates normal heart tissue, while dull fluorescence represents the myocardial infarction area. H: Determination of Myocardial Infarction Area by Fluorescence Section (Fig. 4H). I: Evans blue-TTC Staining Diagram: Red areas depict ischemic but viable myocardial tissue, white areas represent infarcted tissue, and blue areas indicate normal myocardial tissue. J: Myocardial Infarction Area corresponding to Fig. 4I. K: Masson Staining of Frozen Mouse Heart Slices (Scale: 500 μm). L: Degree of Fibrosis corresponding to Fig. 4K. ^&&&&^*P*<0.0001 vs Before MI group; \**P*<0.05, \*\**P*<0.01, \*\*\**P*<0.001, \*\*\*\**P*<0.0001 vs MI group; ^#^*P*<0.05, ^##^*P*<0.01, ^###^*P*<0.001, ^####^*P*<0.0001 vs LncRNA-CIR6 group. (means ± SE, n = 3)

Ultrasonic evaluation revealed a significant increase in postoperative LVIDs and LVIDd, while LVEF and LVFS were significantly lower in the MI group (Fig. 4B). In contrast, the LncRNA-CIR6 and LncRNA-CIR6 + hUCMSCs groups exhibited improvements, with decreased LVIDs and LVIDd, and increased LVEF and LVFS.

Electrical mapping indicated that both the LncRNA-CIR6 group and LncRNA-CIR6 + hUCMSCs group exhibited reduced conduction time and dispersion values and increased conduction velocity compared to the MI group, indicating improved conduction capacity (Fig. 4C-F). Moreover, the combined treatment of LncRNA-CIR6 with hUCMSCs demonstrated superior effects on cardiac contractile function and transmission function after myocardial infarction compared to individual treatments (Fig. 4A-F).

To validate the protective effect of LncRNA-CIR6 or combined hUMSCs treatment on the heart, we analyzed the infarct size and fibrotic surface of postoperative mouse hearts using fluorescence, Evans blue-TTC staining, and Masson staining. Fluorescence results demonstrated that, compared with the MI group, the infarct size of mice in the LncRNA-CIR6 group and LncRNA-CIR6 + hUCMSCs group was significantly reduced (Fig. 4G-H). This finding was consistent with the results of Evans blue-TTC staining (Fig. 4I-J). Masson staining revealed a significant reduction in the area of myocardial fibrosis in both the LncRNA-CIR6 group and LncRNA-CIR6 + hUCMSCs group (Fig. 4K-I). Importantly, the combination of LncRNA-CIR6 with hUMSCs significantly inhibited myocardial infarction size and fibrosis degree compared to treatment alone (Fig. 4G-L). In conclusion, the combination of LncRNA-CIR6 with hUMSCs improves myocardial contractile function and transmission function after infarction *in vivo* and mitigates infarction size and fibrosis.

### 3.3 Prediction of LncRNA-CIR6 Secondary Structure and Its Protein Interactions

To further explore the mechanism of LncRNA-CIR6 in the induction of MSCs, we conducted a biological information analysis. Using Annolnc2, we identified that the transcriptional expression region of LncRNA-CIR6 is on the reverse chain of chromosome 17 in the human genome. The structure of this transcript is very similar to that of the known transcript N-sulfonylglucose sulfate hydrolase(Fig. 5A). Subsequently, we obtained the secondary structure (Fig. 5B) of LncRNA-CIR6 from the database analysis and the mountain chart predicted by the database (Fig. 5C). We interpreted it based on the free energy value of LncRNA molecules, where a lower free energy value indicates higher stability and functionality, making them more likely to be related to diseases. Therefore, based on the mountain chart of LncRNA-CIR6 (Fig. 5C), we predicted that the functional region spanned nucleotides 0-50/190-255 (the area highlighted in red in Fig. 5B).

**Figure 5.**
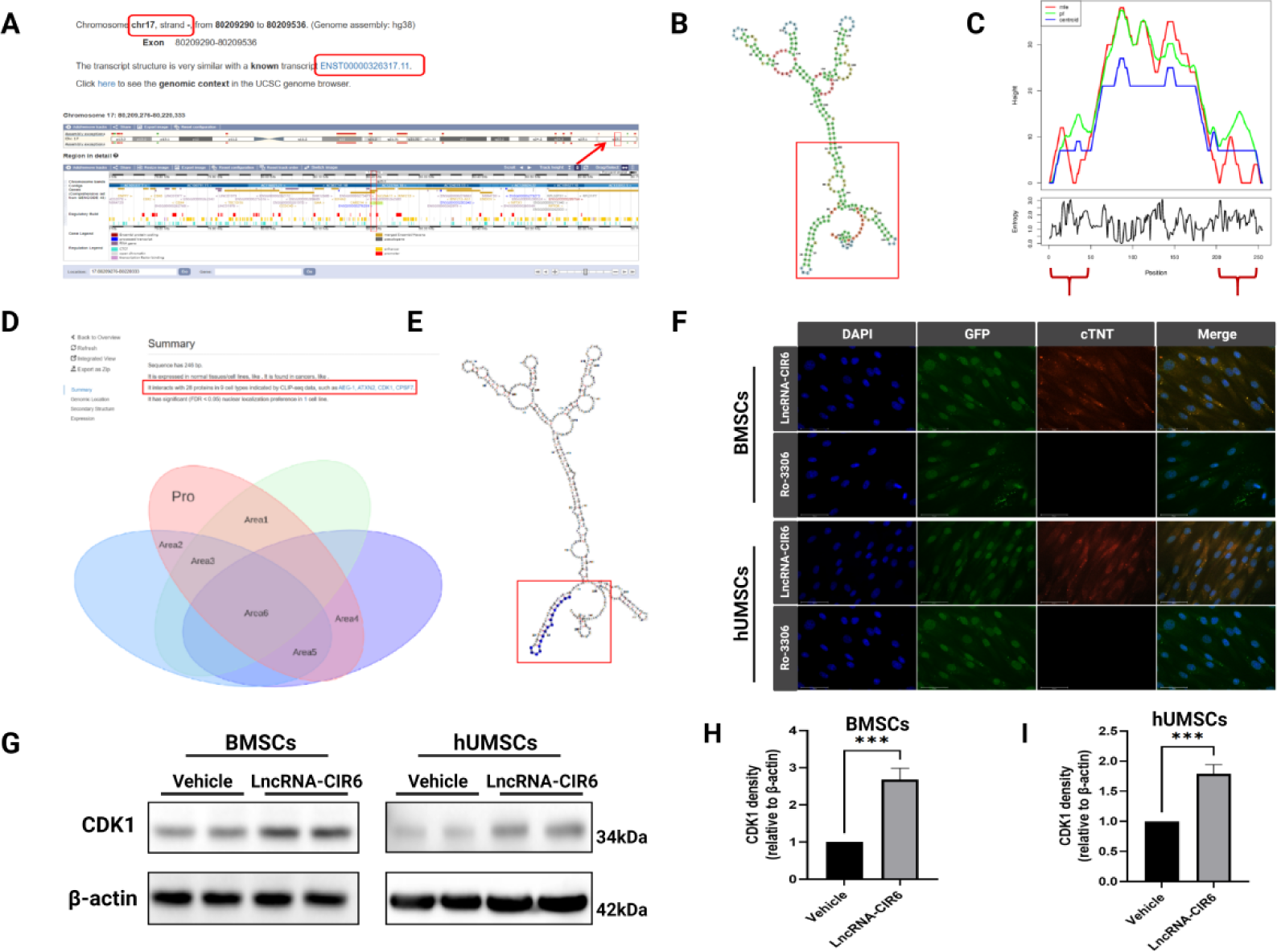
Prediction of LncRNA-CIR6 Secondary Structure and Its Protein Interactions. A: Transcription gene location of LncRNA-CIR6. B: Predicted LncRNA-CIR6 secondary structure. C: Mountain chart of LncRNA-CIR6: LncRNA-CIR6 MFE (minimum free energy) structure, thermodynamic set of RNA structures, and centroid structure. D: Bioinformatics analysis of CDK1. E: CDK1 binding site. F: Immunofluorescence of MSCs (Scale: 20 μm; means ± SE, n = 3). G, H, and I: Western blotting data of CDK1 protein level in MSCs; \*\*\**P*<0.001 vs Vehicle group. (means ± SE, n = 4).

The CLIP-seq data revealed that LncRNA-CIR6 interacts with 28 proteins across nine cell types. Of the 28 predicted proteins, 6 predicted proteins had binding sites larger than 10, were situated in the functional region of LncRNA-CIR6, and were associated with cardiovascular system diseases (Fig. 5D). Notably, CDK1 was the sole protein linked to the proliferation and differentiation of cardiomyocytes [20]. In addition, the binding site of CDK1 in LncRNA-CIR6 was located between 0 and 17 of LncRNA-CIR6, which belonged to the stable functional region (The area highlighted in red box in Fig. 5E). In order to further predict whether CDK1 is involved in LncRNA-CIR6 induction of MSCs, immunofluorescence analysis was performed after induction of MSCs with inhibitors before transfection. Observations revealed that MSCs in the Ro-3306 group displayed no fluorescence in cTNT, whereas the LncRNA-CIR6 group exhibited red fluorescence (Fig. 5F).

Furthermore, CDK1 exhibited high expression in the LncRNA-CIR6 transfection group in Western blot experiments (Fig. 5G), which further confirmed our previous prediction of the association between LncRNA-CIR6 and CDK1. Therefore, we can preliminarily speculate that CDK1 may be involved in the induction of MSCs by LncRNA-CIR6.

## 4. Discussion

Long non-coding RNAs (LncRNAs) have been confirmed to play crucial roles in stem cell pluripotency and cardiac differentiation [21]. LncRNAs constitute an essential part of the transcriptional network in stem cells [22]. Among them, the knockdown or overexpression of LncRNAs can reciprocally influence pluripotency transcription factors, thereby regulating the pluripotent state and lineage specificity of stem cells [23]. Research has identified LncDACH1 as a negative regulator of cardiac regeneration[24]. Its silencing enhances the proliferative potential of myocardial cells, reduces infarct size, and improves cardiac function [24]. Additionally, LncRNAs play a critical role in the process of cellular reprogramming, especially in somatic cell reprogramming [25]. For instance, LncRNA-Bvht is a known LncRNA associated with heart formation during embryonic development, and its expression can prompt embryonic stem cells (ESCs) to differentiate into cardiac muscle cells [26]. These findings provide new insights into the regulatory role of LncRNAs in the proliferation and differentiation of cardiac muscle cells, offering potential therapeutic strategies for research in cardiac regenerative medicine.

LncRNA-CIR6 is a long non-coding RNA specifically expressed in the human fetal heart [27]. Its crucial role is evident in inducing pluripotent stem cells to form cardiomyocytes in both mouse embryos and humans [27]. In addition to expressing markers like cardiac troponin T (cTNT), LncRNA-CIR6 also exhibits cardiac-specific contractile protein markers, including original myosin and α-myosin [27]. Notably, LncRNA-CIR6 possesses an evolutionarily conserved secondary structure, enabling it to promote the differentiation of non-muscle cells into cardiomyocytes [27]. Consequently, we successfully induced the differentiation of MSCs into cardiogenic cells through the transfection of LncRNA-CIR6. The induction of LncRNA-CIR6 significantly improved the efficiency of cardiogenic cell differentiation in MSCs, leading to the induction of the cardiac-specific marker cTNT. These findings suggest that LncRNA-CIR6 may play a crucial role in signaling pathways associated with cardiogenic cell differentiation, thereby promoting the differentiation of MSCs into cardiogenic cells. Despite the currently differentiated cardiogenic cells not displaying beating activity, the indispensable role of LncRNA-CIR6 *in vitro* in inducing MSCs differentiation should not be overlooked. However, the specific mechanism governing this process remains unknown and requires further exploration in subsequent studies.

Existing research has emphasized the potential of stem cell therapy to compensate for damaged cardiac muscle cells, highlighting the critical role of cardiac cell differentiation in replacing those affected by myocardial infarction [28]. *In vitro*, we substantiated the effectiveness of LncRNA-CIR6 in promoting MSC differentiation into cardiogenic cells. In murine models, we observed the positive impact of LncRNA-CIR6 in resisting MI. Particularly promising is the enhanced performance in heart function and reduction in infarct size when LncRNA-CIR6 is combined with hUCMSCs, surpassing the outcomes of using LncRNA-CIR6 alone. This suggests that MSCs transfected with LncRNA-CIR6 may possess superior myocardial regeneration capabilities. Our hypothesis posits that LncRNA-CIR6 may induce the differentiation of endogenous stem cells in the heart or the injected hUCMSCs into cardiogenic cells. Furthermore, studies have demonstrated that cardiac-inducing RNA-6 (CIR-6) can induce the direct differentiation of murine fibroblasts into cardiomyocytes *in vitro* [29]. Therefore, we hypothesize that cardiogenic cells not only derive from the differentiation of MSCs induced by LncRNA-CIR6 but may also transform from myocardial fibroblasts under specific conditions. These regenerated cardiogenic cells play a role in resisting myocardial infarction, offering a potential pathway for heart regeneration. To validate these observations and speculations, further research is planned.

Additionally, we hypothesize that, besides their involvement in the differentiation into cardiogenic cells to replace infarcted cells, both LncRNA-CIR6 and MSCs play distinct roles in conferring cardioprotective effects. In recent years, LncRNAs have garnered significant attention due to their diverse functions and regulatory roles in cellular processes [30]. For instance, LncRNA-NRF has been identified as a regulator of cardiomyocyte death by targeting Mir-873 and RIPK1/RIPK3. The involvement of MSC-derived LncRNAs and exosomes in cardiac injury and repair is well-established [31]. Li et al. elucidated the protective effect of bone marrow mesenchymal stem cell-derived exosomes on hypoxic-reperfusion injury in cardiomyoblasts through the HAND2-AS1/miR-17-5p/Mfn2 axis. Consequently, the possibility that MSC exosomes contribute to heart protection cannot be overlooked. Hence, the investigation into LncRNA-mediated regulation of cardiomyocyte regeneration and proliferation, aimed at enhancing cardiac function post-infarction, warrants further exploration.

In our bioinformatics database analysis, we pinpointed the specific location of the LncRNA-CIR6 transcript and noted its structural similarity to the transcript of n-sulfonyl glucose sulfate hydrolase. The database also predicted proteins interacting with LncRNA-CIR6, with CDK1 being one of them. CDK1 is a protein associated with cardiovascular disease and is involved in the proliferation and differentiation of cardiomyocytes [20]. Under the induction of LncRNA-CIR6, we observed an increased expression of CDK1 in MSCs. Moreover, inhibiting CDK1 made it challenging for MSCs to differentiate into cardiomyocytes. This suggests that CDK1 may play a crucial role in the induction of cardiomyocyte differentiation by LncRNA-CIR6. CDK1, a cyclin-dependent kinase crucial for cell cycle regulation [32], orchestrates cell cycle progression by phosphorylating various substrates [33]. Within the Wnt signaling pathway, CDK1 exerts inhibitory effects through diverse mechanisms [34, 35]. For instance, it phosphorylates LRP6, inhibiting its activity and preventing Wnt ligand binding and signal transduction [35]. Additionally, CDK1 can regulating the Wnt signaling pathway by phosphorylating β-catenin and other components [36].

Studies by Gao et al. propose that inhibiting β-catenin expression facilitates the cardiac differentiation of MSCs [37]. Zhang et al. observed that overexpressing mir-499 in rat BM-MSCs promotes heart-specific gene expression and activates the Wnt/β-catenin signaling pathway by altering the phosphorylated/dephosphorylated β-catenin ratio [38]. Furthermore, some research suggests that Wnt signaling pathway inhibitors enhance the differentiation of mesenchymal stem cells into cardiac progenitors *in vitro*. Inhibiting Wnt signaling also promotes the proliferation of human mesenchymal stem cells, improving cardiomyopathy *in vivo* [39, 40]. Therefore, we hypothesize that LncRNA-CIR6 induces MSC differentiation into cardiogenic cells by regulating the Wnt signaling pathway through CDK1. However, there is still much to unravel regarding the precise mechanisms through which CDK1 regulates the Wnt signaling pathway and influences cardiomyocyte differentiation in MSCs. Further studies are essential to deepen our understanding of these intricate processes.

## Acknowledgments

This work was supported by a Key projects of science and technology development plan of Jilin province (20210204152YY) and National Natural Science Foundation of China (NSFC) grant (82360065)

